# Predicting One-Year Outcome in First Episode Psychosis using Machine Learning

**DOI:** 10.1101/390096

**Authors:** SP Leighton, R Krishnadas, K Chung, A Blair, S Brown, S Clark, K Sowerbutts, M Schwannauer, J Cavanagh, AI Gumley

**Affiliations:** Dr Samuel P Leighton, MBChB, Institute of Health and Wellbeing, University of Glasgow; Dr Rajeev Krishnadas, PhD, Institute of Neuroscience and Psychology, University of Glasgow; Ms Kelly Chung, BSc, Mental Health & Wellbeing, University of Glasgow; Dr Alison Blair, MBChB, ESTEEM First Episode Psychosis Service, NHS Greater Glasgow and Clyde; Dr Susie Brown, MBChB, ESTEEM First Episode Psychosis Service, NHS Greater Glasgow and Clyde; DrSuzy Clark, DCIinPsychol, ESTEEM First Episode Psychosis Service, NHS Greater Glasgow and Clyde; Dr Kathryn Sowerbutts, MBChB, ESTEEM First Episode Psychosis Service, NHS Greater Glasgow and Clyde; Prof Matthias Schwannauer, PhD, Department of Clinical & Health Psychology, University of Edinburgh; Prof Jonathan Cavanagh, MD, Institute of Health and Wellbeing, University of Glasgow; Prof Andrew I Gumley, PhD, Institute of Health and Wellbeing, University of Glasgow

**Author notes:** Joint senior author (J Cavanagh, AI Gumley). **Corresponding Author:** Dr Rajeev Krishnadas, PhD, Honorary Clinical Senior Lecturer, Institute of Neuroscience and Psychology, University of Glasgow.

## Abstract

**Lay Summary:** *Evidence before this study:* Our knowledge of factors which predict outcome in first episode psychosis (FEP) is incomplete. Poor premorbid adjustment, history of developmental disorder, symptom severity at baseline and duration of untreated psychosis are the most replicated predictors of poor clinical, functional, cognitive, and biological outcomes. Yet, such group level differences are not always replicated in individuals, nor can observational results be clearly equated with causation. Advanced machine learning techniques have potential to revolutionise medicine by looking at causation and the prediction of individual patient outcome. Within psychiatry, Koutsouleris *et al* employed machine learning to predict 4- and 52-week functional outcome in FEP to a 75% and 73.8% test-fold balanced accuracy on repeated nested internal cross-validation. The authors suggest that before employing a machine learning model “into real-world care, replication is needed in external first episode samples”.

*Added value of this study:* We believe our study to be the first externally validated evidence, in a temporally and geographically independent cohort, for predictive modelling in FEP at an individual patient level. Our results demonstrate the ability to predict both symptom remission and functioning (in employment, education or training (EET)) at one-year. The performance of our EET model was particularly robust, with an ability to accurately predict the one-year EET outcome in more than 85% of patients. Regularised regression results in sparse models which are uniquely interpretable and identify meaningful predictors of recovery including specific individual PANSS items, and social support. This builds on existing studies of group-level differences and the elegant work of Koutsouleris *et al.*

*Implications of all the available evidence:* We have demonstrated the externally validated ability to accurately predict one-year symptomatic and functional status in individual patients with FEP. External validation in a plausibly related temporally and geographically distinct population assesses model transportability to an untested situation rather than simply reproducibility alone. We propose that our results represent important and exciting progress in unlocking the potential of predictive modelling in psychiatric illness. The next step prior to implementation into routine clinical practice would be to establish whether, by the accurate identification of individuals who will have poor outcomes, we can meaningful intervene to improve their prognosis.

**Abstract:** *Background:* Early illness course correlates with long-term outcome in psychosis. Accurate prediction could allow more focused intervention. Earlier intervention corresponds to significantly better symptomatic and functional outcomes. We use routinely collected baseline demographic and clinical characteristics to predict employment, education or training (EET) status, and symptom remission in patients with first episode psychosis (FEP) at 1 year.

*Methods:* 83 FEP patients were recruited from National Health Service (NHS) Glasgow between 2011 and 2014 to a 24-month prospective cohort study with regular assessment of demographic and psychometric measures. An external independent cohort of 79 FEP patients were recruited from NHS Glasgow and Edinburgh during a 12-month study between 2006 and 2009. Elastic net regularised logistic regression models were built to predict binary EET status, period and point remission outcomes at 1 year on 83 Glasgow patients (training dataset). Models were externally validated on an independent dataset of 79 patients from Glasgow and Edinburgh (validation dataset). Only baseline predictors shared across both cohorts were made available for model training and validation.

*Outcomes:* After excluding participants with missing outcomes, models were built on the training dataset for EET status, period and point remission outcomes and externally validated on the validation dataset. Models predicted EET status, period and point remission with ROC area under curve (AUC) performances of 0.876 (95%CI: 0.864, 0.887), 0.630 (95%CI: 0.612, 0.647) and 0.652 (95%CI: 0.635, 0.670) respectively. Positive predictors of EET included baseline EET and living with spouse/children. Negative predictors included higher PANSS suspiciousness, hostility and delusions scores. Positive predictors for symptom remission included living with spouse/children, and affective symptoms on the Positive and Negative Syndrome Scale (PANSS). Negative predictors of remission included passive social withdrawal symptoms on PANSS.

*Interpretation:* Using advanced statistical machine learning techniques, we provide the first externally validated evidence for the ability to predict 1-year EET status and symptom remission in FEP patients.

*Funding:* The authors acknowledge financial support from NHS Research Scotland, the Chief Scientist Office, the Wellcome Trust, and the Scottish Mental Health Research Network.

## Introduction

Initial clinical presentation and early illness course correlates with long term outcome in first episode psychosis (FEP).^1^ The ability to accurately predict outcome at an individual level would allow more focussed intervention. For patients, a meaningful outcome is often more than simple symptom remission but optimising developmental pathways in early psychosis including vocational and educational outcomes.^(2–4^

FEP most often presents at the critical stage in a young person’s life when community, societal roles, educational and vocational achievement are being forged. Consequently, its onset triggers a precipitous decline in education and employment. Missing out on such vocational opportunities, as enshrined in Article 23 of the Universal Declaration of Human Rights^5^, impairs not only financial independence but also societal inclusion, the forming of relationships and self-actualisation.^6^

Those with FEP want to work. Evidence suggests that finding employment is more important than any specific mental health intervention.^7^ Yet less than a quarter of those with severe mental illness such as schizophrenia receive vocational rehabilitation.^8,9^ Clinicians’ attitudes towards their patients’ ability to return to work and their associated estimation of risks are often ambivalent; this is reinforced by a continued decline in employment rates in the years following contact with mental health services.^10^

Attitudes are changing with a new focus on employment, education and training (EET) recognised by the Meaningful Lives international consensus statement for FEP.^11^ Successful intervention strategies exist, such as the Individual Placement and Support approach in FEP, which have been evidenced to show employment and education rates of 69% as compared with 35% for controls.^10^ If we can correctly identify those with poor EET outcomes at their initial presentation, we could apply such vocational interventions at an earlier stage. Existing evidence suggests this would have a much greater chance of success.^12,13^

At present, our knowledge of factors, which predict outcome in FEP is incomplete. A recent metaanalysis by Lally *et al* provides the first robust evidence of remission and recovery outcomes in FEP but was unable to establish the key clinical or demographic factors, which discriminated between patients. Specifically, it did not replicate an earlier meta-analysis showing an association between longer duration of untreated psychosis (DUP) and worse outcome in FEP.^14,15^ A systematic review identified poor premorbid adjustment, history of developmental disorder, greater symptom severity at baseline and longer DUP to be the most replicated predictors of poor clinical, functional, cognitive, and biological outcomes in early onset psychosis.^16^ Within the Scottish population specifically, socioeconomic deprivation and ethnicity have been shown to be risk factors for developing psychosis, while substance misuse, longer first admission, younger age of onset and male gender increased the risk of poor long-term outcome.^17–19^ However, such group level differences cannot be extrapolated to individuals–the ‘ecological fallacy’^20^–nor can observational results be readily equated with causation or accurate predictions.

Advanced machine learning techniques have potential to revolutionise medicine by looking at causation and the prediction of *individual* patient outcome.^21^ Within psychiatry, machine learning has been already applied to the prediction of response to ECT, from baseline MRI structural data, to an accuracy of 78.3% and sensitivity of 100% on internal validation.^22^. Koutsouleris *et al* employed machine learning to predict 4 and 52-week outcome (Global Assessment of Function ≥65) in FEP to a 75% and 73.8% test-fold balanced accuracy on repeated nested internal cross-validation. The authors suggest that before employing a machine learning model “into real-world care, replication is needed in external first episode samples”.^23(p935)^

Because of the practicalities surrounding the requirement to recruit an additional and independent cohort of patients, few studies assess predictive model generalisability via temporal and/or geographical validation. *External* validation in a plausibly related population is a considerably stronger test of predictive models. It assesses model transportability to an untested situation rather than simply reproducibility alone.^24^ As outlined by Koutsouleris *et al*, this is considered essential before applying the predictive model to clinical practice.^23,25^

Our study is based on the hypothesis that it is possible to predict outcome in terms of employment education or training (EET) status or symptom remission at 1 year in FEP using baseline (pertaining to the time around study entrance) demographic and clinical psychometric biomarkers. We are not aware of any study to date, which has employed predictive modelling of EET status in a FEP cohort. We then assess the generalisability of our predictive model on an external cohort of patients.

Additionally, we seek to find the relative importance of the individual variables in contributing to prediction performance. These could have clinical relevance and may inform future research. Focussing on patients with FEP, who have not had extensive psychotherapeutic or pharmacological interventions, helps mitigate against potential confounders.

## Method

### Participants and Study Design

The Compassionate Recovery: Individualised Support in early Psychosis (CR:ISP) study was a 24-month non-randomised prospective study of individuals with FEP. The study received ethical (11/AL/0247) and institutional review board approval (GN11CP130) and all participants gave informed consent. An Integrated Care Pathway for FEP was implemented which facilitated regular routine assessment of demographic and psychometric measures at time 0, month 3, month 6 and month 12.

Recruitment took place in mental health services in the NHS Greater Glasgow & Clyde (NHSGGC) health board between 2011 and 2014. To be considered for inclusion participants had to be: (a) inpatients or out-patients with (b) first presentation to mental health services for psychosis, (c) ICD-10 diagnosis of non-organic psychosis. 83 participants were entered into the study. The CR:ISP patients formed the training dataset.

The validation dataset was formed of 79 FEP patients recruited to an earlier study (1 September 2006 to 31 August 2009), which took place in mental health services in the NHS in Glasgow and in Edinburgh. Demographic and psychometric measures were assessed at time 0, month 6 and month 12. Participants and study design have been described previously (see ***supplemental materials***).^26^

### Predictor and Outcome Measures

Baseline predictors shared across both cohorts were made available for model training and external validation. Demographic predictors included admission to hospital, age, citizenship, educational attainment, ethnicity, gender, household composition, EET at baseline, parental status, relationship status, accommodation, alcohol use, and recreational drug use status. Psychometric clinical predictors included individual PANSS items^27^ and ordinalised depression rating (training cohort used Hospital Anxiety and Depression Scale (HADS), validation cohort used Beck’s Depression Inventory II (BDI-II)–scores were categorised as none, mild, moderate or severe according to published cut offs.^28, 29^

The 1-year binary outcome measures included EET status at 1-year, PANSS point remission (meeting Andreasen PANSS criteria at month 12), and period remission (meeting Andreasen PANSS criteria at both month 6 and month 12). Andreasen *et al* defined remission as scores of less than or equal to 3 in PANSS items P1 Delusions, P2 Conceptual Disorganisation, P3 Hallucinatory Behaviour, N1 Blunted Affect, N4 Apathetic Social Withdrawal, N6 Lack of Spontaneity and G9 Unusual Thought Content, present for a period of at least 6 months.^30^

### Statistical Analysis

All statistical analyses were carried out within the R programming environment.^31^

Between-group (training versus validation) differences were tested using Welch’s independent t-test for continuous variables and Pearson’s chi-squared or Fisher’s exact test for categorical variables. Bonferroni correction was employed for multiple comparisons.

Machine learning analysis was carried out using the ‘Caret’ package.^32^ R code is available in the supplemental materials.

During pre-processing, data were centred and scaled, variables with zero variance and near-zero variance removed and missing data imputed using k (5) nearest neighbour imputation, prior to model construction. In psychiatric research, data are commonly seen as ‘missing not at random’. For example, with drop-out more frequent in those who have relapsed. Ignoring this leads to systematic attrition bias in any inference drawn, in addition to a loss of power. Multiple imputation allows for nearly unbiased parameter estimates.^33,34^

A logistic regression model was fit by elastic net regularisation with variable selection in ‘Caret’ with the ‘glmne package.^35^ ‘Glmnet’ fits a generalized linear model via penalised maximum likelihood. The objective function for the penalised logistic regression uses the negative binomial log-likelihood:

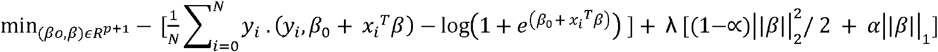

The alpha (α) hyperparameter determines the balance between ridge (α = 0) and lasso (α = 1) penalty and the lambda (*λ*) hyperparameter determines the amount of penalty. Elastic net regression improves upon ordinary least squares via regularisation (shrinkage) of the estimated beta coefficients (β). This results in superior performance when number of predictors (p) approaches the number of subjects (N) (in such circumstances there is no unique least squares estimate), or in the presence multicollinearity by offsetting a small amount of bias with large reductions in variance. Further, a consequence of regularisation is feature selection, which improves model interpretability.^36^

We used n(5) fold repeated (100 times) cross validation to train and tune our model over a grid of alpha and lambda hyperparameters on our training dataset (see supplemental materials). All splits were balanced by outcome class. The best model was refit on the whole training set to calculate its standardised beta coefficients. The exponentials of which, the odds ratios, are presented.

We estimated the discriminative performance of the models using receiver operating curve (ROC) area under the curve (AUC). ROCAUC is independent of class distribution and selects for models that achieve false positive and true positive rates that are significantly above random chance.

External validity was established by using the model built on the training dataset to predict the probability of the outcome class and comparing it to the actual outcome class in the validation dataset, then, calculating the ROC AUC performance metric using the ‘pROC’ package.^37,38^ The 95%CI of the ROCAUC was established based on U-statistic theory using the ‘clinfun’ package. ^38,39^ In addition, permutation testing was used to confirm significance, whereby, the actual ROCAUC was compared to its null ROCAUC distribution derived by testing the model on randomly permutated class outcomes, repeated 10000 times. The p value is the proportion of permutated values greater than or equal to the actual value.^40^ The model accuracy, sensitivity, specificity, positive predictive value (PPV) and negative predictive value (NPV) are presented based on the point closest to the top-left of the ROC plot. The formulae for these quantities have been described previously.^41^

The above procedure was repeated for each of the three dependent outcome variables.

## Results

**Table 1** summarises the predictor and outcome data for the training and validation cohorts. The only statistically significantly different predictor variable between the cohorts was accommodation. More people lived in rented accommodation in the training cohort, and more people lived with their family in the validation cohort (p<0.001). There were no statistically significant differences in each of the three outcomes between the cohorts.

**Table 1.** Characteristics of training cohort and validation cohort.

| **BASELINE**<br>**PREDICTOR** | **TRAINING COHORT**<br>**(N=83)** | **MISSING**<br>**NO. (%)** | **VALIDATION**<br>**COHORT (N=79)** | **%**<br>**MISSING** | **P-**<br>**VALUE** |
| --- | --- | --- | --- | --- | --- |
| **ACCOMODATION**<br>**NO. (%)** | Rented – 48 (59)<br>Private with family – 30 (37)<br>Private owner – 4 (5)<br>Homeless – 0 (0)<br>NFA – 0 (0)<br>Prison – 0 (0) | 1 (1) | Rented – 20 (28)<br>Private with family – 43 (61)<br>Private owner – 4 (6)<br>Homeless – 2 (3)<br>NFA – 1 (1)<br>Prison – 1 (1) | 8 (10) | <0.001* |
| **AGE**<br>**MEAN (SD),**<br>**YEARS** | 25.22<br>(19.68, 30.75) | 0 (0) | 24.64<br>(17.46, 31.72) | 3 (4) | 0.6 |
| **ALCOHOL USE**<br>**NO. (%)** | 58 (74) | 5 (6) | 58 (85) | 11 (14) | 0.1 |
| **CITIZENSHIP**<br>**NO. (%)** | UK – 72 (87)<br>Other – 5 (6)<br>Asylum seeker – 4 (5)<br>Refugee – 2 (2) | 0 (0) | UK – 74 (95)<br>Other – 3 (4)<br>Asylum seeker – 0 (0)<br>Refugee – 1 (1) | 1 (1) | 0.2 |
| **DEPRESSION**<br>**RATING**<br>**NO. (%)** | None – 37 (50)<br>Mild – 11 (15)<br>Moderate – 13 (18)<br>Severe – 13 (18) | 9 (11) | None – 16 (32)<br>Mild – 10 (20)<br>Moderate – 11 (22)<br>Severe – 13 (26) | 29 (37) | 0.3 |
| **DRUG USE**<br>**NO. (%)** | 42 (58) | 11 (13) | 37 (64) | 21 (27) | 0.5 |
| **HIGHEST EDUCATIONAL ATTAINMENT**<br>**NO. (%)** | Before 16 – 14 (18)<br>At 16 – 17 (21)<br>17 to 18 – 21 (26)<br>College – 19 (23)<br>University – 9 (11) | 3 (4) | Before 16 – 16 (23)<br>At 16 – 17 (24)<br>17 to 18 – 20 (28)<br>College – 11 (15)<br>University – 7 (10) | 8 (10) | 0.7 |
| **ETHNICITY**<br>**NO. (%)** | White – 67 (81)<br>Other – 16 (19) | 0 (0) | White – 74 (96)<br>Other – 3 (4) | 2 (3) | 0.003 |
| **GENDER**<br>**NO. (%)** | Male – 55 (66)<br>Female – 28 (34) | 0 (0) | Male – 54 (68)<br>Female – 25 (32) | 0 (0) | 0.8 |
| **HOSPITAL ADMISSION AT BASELINE**<br>**NO. (%)** | 32 (39) | 1 (1) | 40 (51) | 1 (1) | 0.1 |
| **OTHERS IN HOUSEHOLD**<br>**NO. (%)** | Parents & siblings – 42 (51)<br>Alone – 15 (18)<br>Spouse & children – 13 (16)<br>Friends – 9 (11)<br>Siblings – 3 (4) | 1 (1) | Parents & siblings – 38 (55)<br>Alone – 22 (32)<br>Spouse & children – 5 (7)<br>Friends – 4 (6)<br>Siblings – 0 (0) | 10 (13) | 0.06 |
| **PANSS TOTAL**<br>**MEAN (SD)** | 83.78<br>(59.16, 108.4) | 1 (1) | 74.43<br>(52.93, 95.94) | 3 (4) | 0.01 |
| **PARENT**<br>**NO. (%)** | 13 (16) | 0 (0) | 9 (12) | 1 (1) | 0.4 |
| **EMPLOYMENT,**<br>**EDUCATION OR**<br>**TRAINING**<br>**NO. (%)** | 45 (56) | 2 (2) | 40 (52) | 2 (3) | 0.6 |
| **RELATIONSHIP**<br>**STATUS** | Single – 65 (78)<br>Relationship –<br>12 (14)<br>Married – 3 (4)<br>Separated – 3 (4)<br>Divorced – 0 (0) | 0 (0) | Single – 69 (88)<br>Relationship –<br>5 (6)<br>Married – 1 (1)<br>Separated – 2 (2)<br>Divorced – 1 (1) | 1 (1) | 0.3 |
| **ONE-YEAR**<br>**OUTCOME** | **TRAINING COHORT**<br>**NO. (%)** |  | **VALIDATION**<br>**COHORT NO. (%)** | **P VALUE** |  |
| **PERIOD**<br>**REMISSION** | 33 / 67<br>(49) |  | 33 / 64<br>(52) | 0.8 |  |
| **POINT**<br>**REMISSION** | 40 / 71<br>(56) |  | 46 / 67<br>(69) | 0.1 |  |
| **EMPLOYMENT,**<br>**EDUCATION OR**<br>**TRAINING** | 32 / 75<br>(43) |  | 27 / 67<br>(40) | 0.8 |  |

\* indicates Bonferroni corrected significance ( $p = 2.78 \times 10^{-3}$ ). SD = standard deviation; NO. = number of subjects.

**Table 2** summarises the EET status, point remission, and period remission models’ externally validated performance. **Figure 1** show each model’s respective ROC curve–EET status **(A)**, point remission **(B)** and period remission **(C).** All three models’ ROCAUC and 95%CI were significantly better than chance. The model predicting EET status had particularly high performance with a ROCAUC of 0.876 (95%CI: 0.864, 0.887; p = <0.001).

**Table 2.**
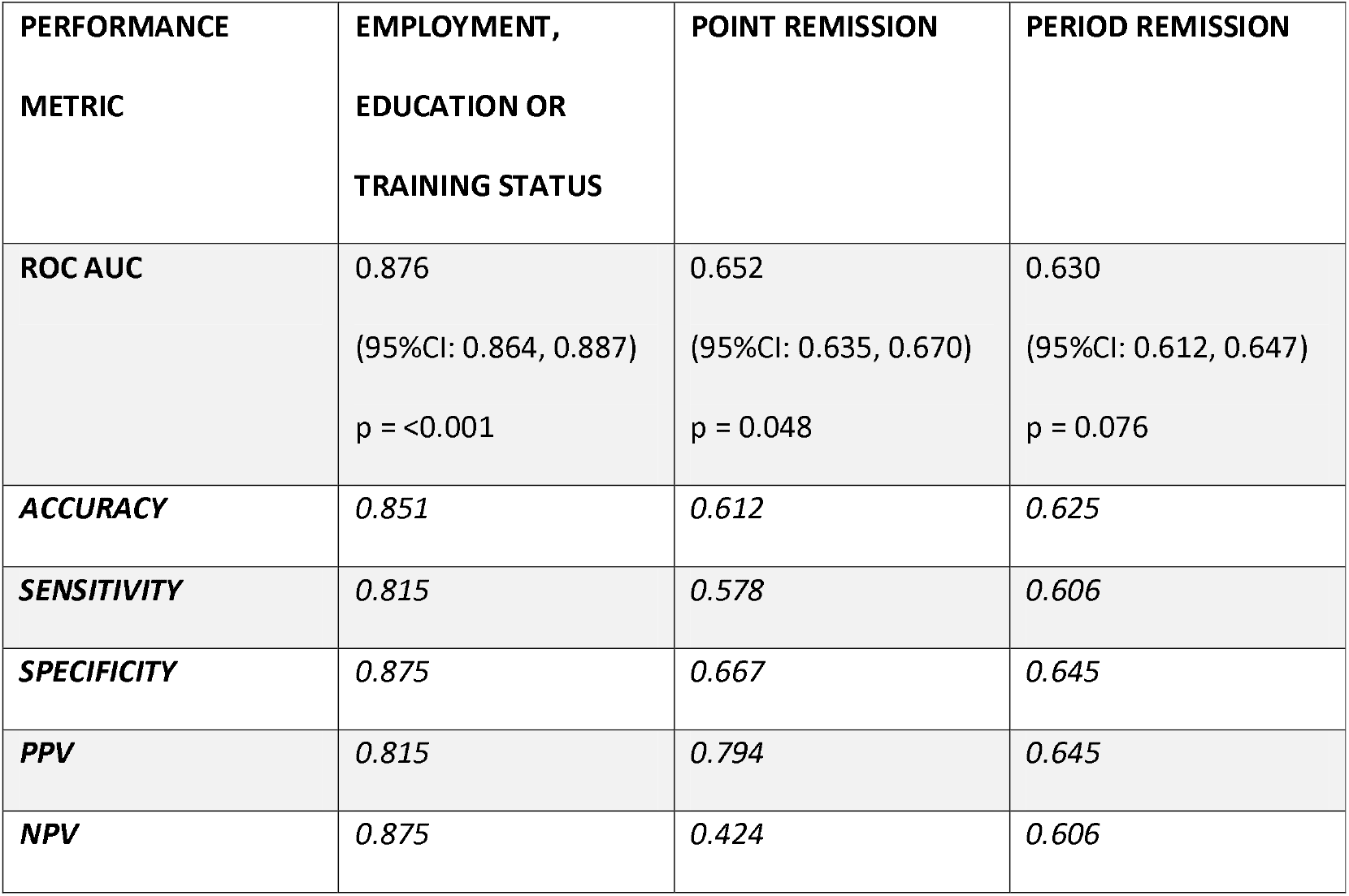
One-year EET status, point remission, and period remission models’ externally validated performance metrics. The 95%CI of the ROC AUC was established based on U-statistic theory, and significance level by permutation testing (n = 10000). Representative accuracy, sensitivity, specificity, positive predictive, and negative predictive values are presented in italics based on the point on the ROC curve closest to the top left. ROC AUC = Receiver Operating Characteristic Area Under the Curve; PPV = Positive Predictive Value; NPV = Negative Predictive Value; 95%CI = 95% Confidence Interval.

**Figure 1.**
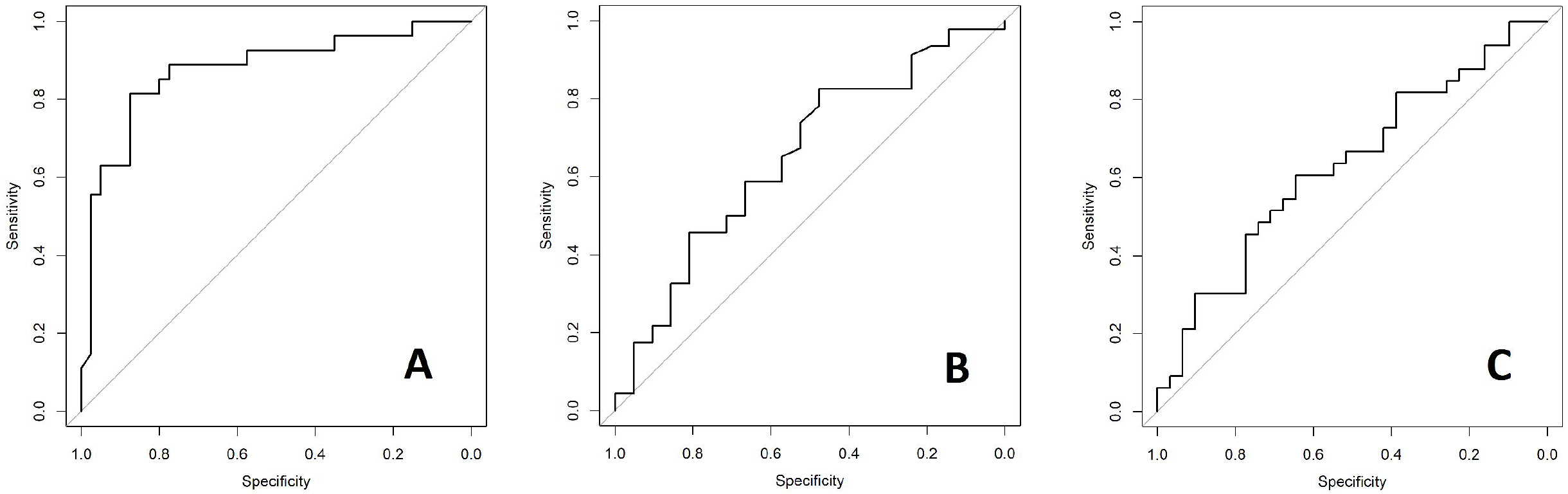
ROC curves for model of one-year EET status **(A),** point remission **(B),** and period remission **(C).** ROC AUC = Receiver Operating Characteristic Area Under the Curve; EET = Employment, Education or Training.

**Figures 2 and 3** summarise the feature selected predictors and their odds ratios for the EET and point remission models. Those for the period remission model are summarised in **Supplemental Figure 1.**

**Figure 2.**
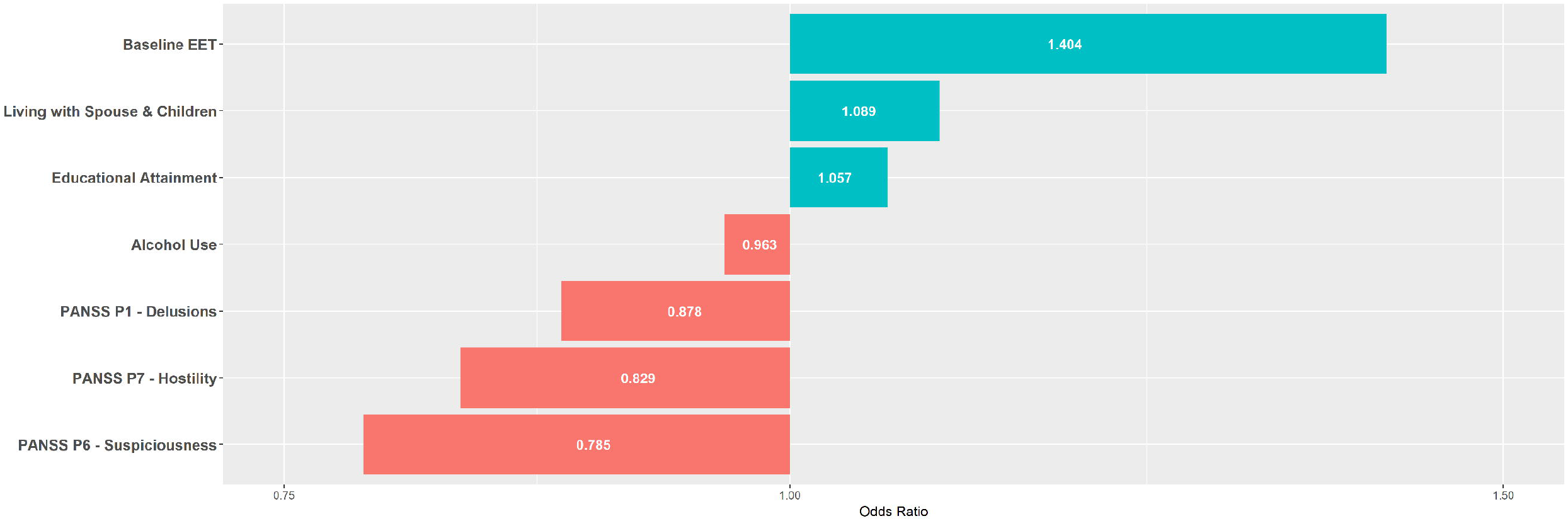
One-year EET status model’s predictors and respective odds ratios. Features selected by elastic net regularisation and respective odds ratios for one-year EET status model. EET = Employment, Education or Training; PANSS = Positive And Negative Symptoms Scale.

**Figure 3.**
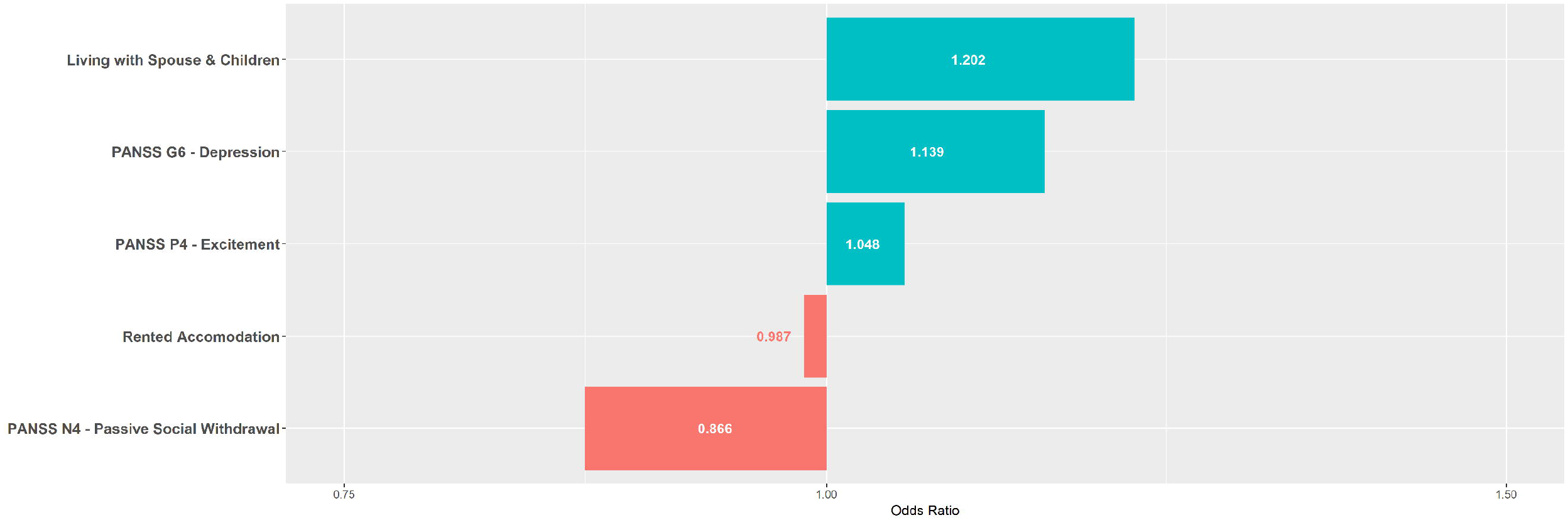
One-year point remission model’s predictors and respective odds ratios. Features selected by elastic net regularisation and respective odds ratios for one-year point remission model. PANSS = Positive And Negative Symptoms Scale.

## Discussion

We believe our study to be the first externally validated evidence, in a temporally and geographically independent cohort, for predictive modelling in FEP at an individual patient level. This builds on existing studies of group-level differences and the elegant work of Koutsouleris *et al*, which outlined the first internally validated evidence of the ability to predict functional outcome in individual patients with FEP.^23^

Our results demonstrate the ability to predict both symptom remission and more importantly, functioning (in employment, education or training). The performance of our EET model was particularly robust, with an ability to accurately predict the one-year EET outcome in more than 85% of patients. This is clinically relevant, especially in the context of the 2010 international first episode psychosis “Meaningful Lives” consensus statement on unemployment as a source of significant social disability.^11^

Predictive statistics focus on the predictive performance of models on new ‘unseen’ data. This helps avoid the inherent problems with conventional statistical techniques, which only explore relationships within the study dataset. Few such relationships are externally validated and, of those that are, many lose significance.^42,43^

Because of the inherent feature selection of regularised logistic regression classifiers, which avoids overfitting in ill-conditioned regression problems where the number of variables is close to the sample size, our models are sparse and uniquely interpretable. Further, without *a priori* selection of variables but instead allowing the classifier to select features, we avoid the introduction of additional observer bias.^35,36^

The period remission model shares all the same predictors that are selected for the point remission model. However, the period remission model selects for additional predictors but with a reduced performance. This suggests possible overfitting on the training dataset, whereby the signal to noise variable ratio captured by the model is reduced, hence poorer generalisability to the independent validation dataset.^44^

Focussing on the shared predictors for both point and period remission, higher scores on the individual PANSS items P4 Excitement and G6 Depression correspond to a positive outcome. Over a century later, this continues to lend credence to a central tenant of the Kraepelian dichotomy–that psychosis in the context of phenomenologically prominent affective presentations tends to have a better chance of recovery.^45^ Recent research in this area is conflicting and hampered by the few prospective studies looking at the influence of affective symptoms on outcome.^46^

PANSS item N4 passive or apathetic social withdrawal predicts poor one-year remission outcomes, a so called negative symptom frequently found in the classical schizophrenia syndrome.^47^ We have previously identified social and interpersonal factors as influencing outcome in FEP.^26,48^ Such initial negative symptoms persist longer and are more difficult to treat, with limited evidence for benefit with conventional pharmacotherapy.^49^ In contrast, there is better evidence in support of psychosocial interventions. For example, cognitive behavioural therapy has been shown to be effective up to 24-months.^50^ Recent meta-analysis supports the use of interventions which enhance social cognition and interpersonal skills in the treatment of negative symptoms in psychosis.^51^ Our finding highlights the requirement for a multimodal approach to treatment of FEP earlier in the illness course. Further, our model selects variables representative of higher social support including living with spouse and children and not renting as predictive of good symptomatic outcome which are consistent with established literature.^52^

Similarly, living with spouse and children is predictive of a good one-year EET outcome. Again, consistent with established literature, our model selects for good baseline EET and higher educational attainment as positive predictors of positive one-year EET outcome.^52^ Interestingly, different individual PANSS items–P1 Delusions, P6 Suspiciousness and P7 Hostility–emerged as negative predictors of one-year EET. Such features may hinder service engagement and attachment which is known to influence recovery and adaption in FEP.^26^ That distinct variables predict EET compared to those predicting remission, suggests the outcomes are capturing different yet complementary areas of recovery.

Our focus on EET status is timely and reflects the growing recognition within the literature that this functional outcome is at least as important as symptom remission. Young people not being in employment, education or training (NEET) has a considerable economic cost, accounting for a loss of €153bn or 1.2% of the European Union’s gross domestic product. Prolonged economic inactivity leads to serious mental health sequelae with higher rates of depression, alcohol or substance misuse and increased suicide attempts. Reducing youth unemployment remains a policy priority in high-income countries including the USA and Europe but the quality of evidence available to legislators surrounding NEET status is low.^53^ We suggest that our robust finding related to the prediction of NEET outcome in first episode psychosis is the first high quality externally validated evidence.

The use of an externally (temporally and geographically) validated dataset with the same predictor and outcome variables is a major strength. Further, our models are interpretable, employ easily obtained variables and, especially in relation to prediction of EET status, are robust. However, our study has limitations. Small sample size relative to the number of predictors is a potential concern. Using elastic net regularised regression, which simulation studies have shown outperforms conventional ordinary least squares or lasso while still enjoying the interpretability of feature selection missing with ridge, helps circumvent such concerns. Another concern may be that our cohorts have a statistically significant difference in accommodation status with more participants renting and less living in private accommodation with family in the training cohort as compared with the validation cohort. Similarly, changes in the wider macroeconomic environment, such as those following the 2008 financial crisis which would have been felt by the training cohort only, are known to influence mental health and functional outcomes.^54^ However, we would contend that such differences are in fact a strength of our study and reinforces the importance of external validation. Despite these, and that our model includes renting as a negative predictor of EET status, it still had a statistically significant and robust predictive performance in a geographically and temporally distinct cohort. A further limitation is the lack of biological and additional social variables, such as nicotine dependence^55^, which observational studies have linked to outcomes in psychosis. These should be addressed when designing future studies.

Finally, a required next step prior to implementation into routine clinical practice would be to establish whether, by the accurate identification of individuals who will have poor outcomes, we can meaningful intervene to improve their prognosis.

## Conclusions

We have demonstrated that it is possible to accurately predict one-year symptomatic and functional employment education or training status in first episode psychosis. This has not been reported previously in an externally validated cohort. We propose that our results represent an important and exciting next step in unlocking the potential of predictive modelling in psychiatric illness.

## Acknowledgments

The authors acknowledge financial support from NHS Research Scotland, the Chief Scientist Office, the Wellcome Trust and of the Scottish Mental Health Research Network.

## References

1 Petersen L, Nordentoft M, Jeppesen P, et al. Improving 1-year outcome in first-episode psychosis: OPUS trial. Br J Psychiatry Suppl 2005; 48: s98–103.

2 Lambert M, Schimmelmann BG, Naber D, et al. The Journal of clinical psychiatry. [Physicians Postgraduate Press], 2006 http://www.psychiatrist.com/jcp/article/Pages/2006/v67n11/v67n1104.aspx (accessed Jan 18, 2018).

3 Bond GR, Drake RE, Luciano A. Employment and educational outcomes in early intervention programmes for early psychosis: a systematic review. Epidemiol Psychiatr Sci 2015; 24: 446–57.

4 Tapfumaneyi A, Johnson S, Joyce J, et al. Predictors of vocational activity over the first year in inner-city early intervention in psychosis services. Early Interv Psychiatry 2015; 9: 447–58.

5 United Nations. Universal Declaration of Human Rights. 1948. http://www.ohchr.org/EN/UDHR/Documents/UDHR-Translations/eng.pdf (accessed Jan 18, 2018).

6 Maslow AH. A theory of human motivation. Psychol Rev 1943; 50: 370–96.

7 Shepherd G. Perspectives on schizophrenia: A survey of user, family carer and professional views regarding effective care. J Ment Heal 1995; 4: 403–22.

8 Lehman AF, Steinwachs DM. Patterns of usual care for schizophrenia: initial results from the Schizophrenia Patient Outcomes Research Team (PORT) Client Survey. Schizophr Bull 1998; 24: 11–20–32.

9 Bertram M, Howard L. Employment status and occupational care planning for people using mental health services. Psychiatr Bull 2006; 30: 48–51.

10 Rinaldi M, Killackey E, Smith J, Shepherd G, Singh SP, Craig T. First episode psychosis and employment: A review. Int Rev Psychiatry 2010; 22: 148–62.

11 International First Episode Vocational Recovery (iFEVR) Group. Meaningful lives: Supporting young people with psychosis in education, training and employment: an international consensus statement. Early Interv Psychiatry 2010; 4: 323–6.

12 Killackey EJ, Jackson HJ, Gleeson J, Hickie IB, Mcgorry PD. Exciting Career Opportunity Beckons! Early Intervention and Vocational Rehabilitation in First-Episode Psychosis: Employing Cautious Optimism. Aust New Zeal J Psychiatry 2006; 40: 951–62.

13 Craig T, Shepherd G, Rinaldi M, et al. Vocational rehabilitation in early psychosis: cluster randomised trial. Br J Psychiatry 2014; 205: 145–50.

14 Lally J, Ajnakina O, Stubbs B, et al. Remission and recovery from first-episode psychosis in adults: systematic review and meta-analysis of long-term outcome studies. Br J Psychiatry 2017; 211: 350–8.

15 Marshall M, Lewis S, Lockwood A, Drake R, Jones P, Croudace T. Association Between Duration of Untreated Psychosis and Outcome in Cohorts of First-Episode Patients. Arch Gen Psychiatry 2005; 62: 975.

16 Díaz-Caneja CM, Pina-Camacho L, Rodríguez-Quiroga A, Fraguas D, Parellada M, Arango C. Predictors of outcome in early-onset psychosis: a systematic review. npj Schizophr 2015; 1: 14005.

17 Srireddy P, Agnihotri A, Park J, Taylor J, Connolly M, Krishnadas R. Ethnicity, deprivation and psychosis: the Glasgow experience. Epidemiol Psychiatr Sci 2012; 21: 311–6.

18 Queirazza F, Semple DM, Lawrie SM. Transition to schizophrenia in acute and transient psychotic disorders. Br J Psychiatry 2014; 204: 299–305.

19 Alderson HL, Semple DM, Blayney C, Queirazza F, Chekuri V, Lawrie SM. Risk of transition to schizophrenia following first admission with substance-induced psychotic disorder: a population-based longitudinal cohort study. Psychol Med 2017; 47: 2548–55.

20 Sedgwick P. Understanding the ecological fallacy. BMJ 2015; 351: h4773.

21 Darcy AM, Louie AK, Roberts LW. Machine Learning and the Profession of Medicine. JAMA 2016; 315: 551.

22 Redlich R, Opel N, Grotegerd D, et al. Prediction of Individual Response to Electroconvulsive Therapy via Machine Learning on Structural Magnetic Resonance Imaging Data. JAMA Psychiatry 2016; 73: 557.

23 Koutsouleris N, Kahn RS, Chekroud AM, et al. Multisite prediction of 4-week and 52-week treatment outcomes in patients with first-episode psychosis: a machine learning approach. The Lancet Psychiatry 2016; 3: 935–46.

24 Justice AC, Covinsky KE, Berlin JA. Assessing the generalizability of prognostic information. Ann Intern Med 1999; 130: 515–24.

25 Steyerberg EW, Vergouwe Y. Towards better clinical prediction models: seven steps for development and an ABCD for validation. Eur Heart J 2014; 35: 1925–31.

26 Gumley Al, Schwannauer M, Macbeth A, et al. Insight, duration of untreated psychosis and attachment in first-episode psychosis: prospective study of psychiatric recovery over 12-month follow-up. Br J Psychiatry 2014; 205: 60–7.

27 Kay SR, Fiszbein A, Opler LA. The Positive and Negative Syndrome Scale (PANSS) for Schizophrenia. Schizophr Bull 1987; 13: 261–76.

28 Zigmond AS, Snaith RP. The hospital anxiety and depression scale. Acta Psychiatr Scand 1983; 67:361–70.

29 Beck AT, Steer RA, Brown GK. BDI-II, Beck depression inventory⍰: manual, 2nd ed. San Antonio Tex. ⍰:Boston: Psychological Corp., 1996 http://www.worldcat.org/title/bdi-ii-beck-depression-inventory-manual/oclc/36075838 (accessed Jan 18, 2018).

30 Andreasen NC, Carpenter WT, Kane JM, Lasser RA, Marder SR, Weinberger DR. Remission in Schizophrenia: Proposed Criteria and Rationale for Consensus. Am J Psychiatry 2005; 162: 441–9.

31 R Core Team. R: A language and environment for statistical computing. 2017. https://www.r-project.org/.

32 Kuhn M. caret: Classification and Regression Training. https://github.com/topepo/caret/.

33 von Hippel P, Lynch J. Efficiency Gains from Using Auxiliary Variables in Imputation. 2013; published online Nov 20. http://arxiv.org/abs/1311.5249 (accessed Jan 18, 2018).

34 Sterne JAC, White IR, Carlin JB, et al. Multiple imputation for missing data in epidemiological and clinical research: potential and pitfalls. BMJ 2009; 338: b2393.

35 Friedman J, Hastie T, Tibshirani R. Regularization Paths for Generalized Linear Models via Coordinate Descent. J Stat Softw 2010; 33: 1–22.

36 Zou H, Hastie T. Regularization and variable selection via the elastic net. J R Stat Soc Ser B (Statistical Methodol 2005; 67: 301–20.

37 Robin X, Turck N, Hainard A, et al. pROC: an open-source package for R and S+ to analyze and compare ROC curves. BMC Bioinformatics 2011; 12: 77.

38 DeLong ER, DeLong DM, Clarke-Pearson DL. Comparing the Areas under Two or More Correlated Receiver Operating Characteristic Curves: A Nonparametric Approach. Biometrics 1988; 44: 837.

39 Seshan VE. clinfun: Clinical Trial Design and Data Analysis Functions. 2017. https://cran.r-project.org/package=clinfun.

40 Pérez-Guaita D, Kuligowski J, Garrigues S, Quintás G, Wood BR. Assessment of the statistical significance of classifications in infrared spectroscopy based diagnostic models. Analyst 2015; 140:2422–7.

41 Leighton SP, Nerurkar L, Krishnadas R, Johnman C, Graham GJ, Cavanagh J. Chemokines in depression in health and in inflammatory illness: a systematic review and meta-analysis. Mol Psychiatry 2018; 23: 48–58.

42 Schooler JW. Metascience could rescue the ‘replication crisis’. Nature 2014; 515: 9–9.

43 Open Science Collaboration OS. Estimating the reproducibility of psychological science. Science 2015; 349: aac4716.

44 Waldron L, Pintilie M, Tsao M-S, Shepherd FA, Huttenhower C, Jurisica I. Optimized application of penalized regression methods to diverse genomic data. Bioinformatics 2011; 27:3399–406.

45 Kraepelin E. Psychiatrie: Ein Lehrbuch für Studirende und Aerzte I. Leipzig: J. A. Barth, 1899 https://archive.org/details/psychiatrieeinle02krae (accessed Jan 18, 2018).

46 Jäger M, Haack S, Becker T, Frasch K. Schizoaffective disorder--an ongoing challenge for psychiatric nosology. Eur Psychiatry 2011; 26: 159–65.

47 Boonstra N, Klaassen R, Sytema S, et al. Duration of untreated psychosis and negative symptoms —A systematic review and meta-analysis of individual patient data. Schizophr Res 2012; 142: 12–9.

48 Mcleod H, Gumley A, Macbeth A, Schwannauer M, Lysaker. Metacognitive functioning predicts positive and negative symptoms over 12 months in first episode psychosis. J Psychiatr Res 2014; 54: 109–15.

49 Leucht S, Heres S, Kissling W, Davis JM. Evidence-based pharmacotherapy of schizophrenia. Int J Neuropsychopharmacol 2011; 14: 269–84.

50 National Collaborating Centre for Mental Health (UK). Psychosis and Schizophrenia in Adults; NICE Clinical Guidelines, No. 178. London: National Institute for Health and Care Excellence (UK), 2014 http://www.ncbi.nlm.nih.gov/pubmed/25340235 (accessed Jan 18, 2018).

51 Turner DT, McGlanaghy E, Cuijpers P, van der Gaag M, Karyotaki E, MacBeth A. A Meta-Analysis of Social Skills Training and Related Interventions for Psychosis. Schizophr Bull 2017; published online Nov 11. DOI:10.1093/schbul/sbx146.

52 Rosen K, Garety P. Predicting Recovery From Schizophrenia: A Retrospective Comparison of Characteristics at Onset of People With Single and Multiple Episodes. Schizophr Bull 2005; 31: 735–50.

53 Scott J, Fowler D, McGorry P, et al. Adolescents and young adults who are not in employment, education, or training. BMJ 2013; 347: f5270.

54 Ramanathan S, Balasubramanian N, Krishnadas R. Macroeconomic Environment During Infancy as a Possible Risk Factor for Adolescent Behavioral Problems. JAMA Psychiatry 2013; 70: 218.

55 Krishnadas R, Jauhar S, Telfer S, Shivashankar S, McCreadie RG. Nicotine dependence and illness severity in schizophrenia. Br J Psychiatry 2012; 201: 306–12.

